# Decoding ladybird’s colours: structural mechanisms of colour production and pigment modulation

**DOI:** 10.1101/2025.01.16.633144

**Authors:** Marzia Carrada, Mohamed Haddad, Luis San José, Gonzague Agez, Jean-Marie Poumirol, Alexandra Magro

## Abstract

This study investigates the mechanisms underlying colour production in the family Coccinellidae, focusing on two model species: *A. bipunctata (L)* and *C. quatuordecimguttata (L)*. In this family, colours have traditionally been attributed primarily to pigments such as carotenoids and melanins. We propose an alternative perspective, considering the elytra as an integrated optical medium whose optical properties result from both its architectural design and the properties of its constituent materials, including matrix and pigments. In the present work, several methods were used, ranging from optical and electron microscopies, light-based techniques, to LC-HRMS analyses. Through local pigment analyses, microstructural examination of elytra, and modelling of interaction of elytra with light, we show that, while pigments are central to patterning and contribute to colour, overall colour also results from one or more physical mechanisms that may operate simultaneously. In the light of these results, we assume that the complex and diverse colouration in the Coccinellidae can only be elucidated by considering the interplay of pigments and the optical properties of the elytron cuticle.

## Introduction

Colouration displayed by animals has been the target of intense research aiming at understanding how it is produced, both proximately (*i.e.*, its developmental, genetic and morphological determinants) and ultimately (*i.e.*, its adaptive role and its evolutionary origin and stability) (1). The pigmentary origin of colouration has received a predominant attention probably because of pigments’ inherent properties (namely pleiotropic effects on individuals’ physiology and behaviour) having allowed biologists to link colourful displays to their signalling role in distinct contexts such as reproduction or aposematism (2–4). However, pigments are likely to explain only a small fraction of nature’s colour palette (*e.g.*, (5,6)). Moreover, pigments must rely on underlying, often complexly elaborated, reflective structures to produce any colouration (7,8). Although the structural part of colouration is gaining recognition (*e.g.*, (9–11), its contribution to the production and the signalling function of colour traits is largely neglected, particularly when pigments are also present in the coloured integument. Structural colouration has been less thoroughly studied than pigment-based colouration for different reasons. First, structural components, contrary to pigments, lack a clear framework explaining how they can signal genetic or phenotypic quality. In the same line, the genetic pathways building up structural colourations are poorly known compared to those of pigments (12) even in well-studied taxa such as birds (13). Nevertheless, evidence showing that structural colouration does associate with quality aspects such as individual condition accumulates in different taxa (14). However, the lack of a clear link to the potential costs or constraints that animals face when developing or maintaining structures contributing to colour has limited our understanding of their relevance for the evolution of colourful displays, particularly in animal communication contexts. Secondly, the study of structural colouration requires a good understanding of the physics of light and how small changes in integumentary structures translate into colour changes. Indeed, teguments can be very complex structures that interact with light in many different ways: structural colours can be generated, depending on the tegument’s microstructure, by several physical phenomena such as interference, diffraction, or diffusion, but also by nanostructures such as photonic crystals. Some of these physical mechanisms may occur at the same time and the light interaction with the structure becomes very complicated and hard to describe (15). As a consequence, structural and pigment-based colourations have often been studied in isolation (16), maintaining a long-standing dichotomy that classifies colour traits as either strictly structural or pigment based (17). This established dichotomy contrasts with the fact that it is generally acknowledged that pigments need underlying reflective structures to reveal their light absorption effect and the fact that colouration results from combined more than just additive effects of both structures and pigments.

Insects have been central to the study of animal colouration. However, structural and pigment-based colourations in insects have been most often considered separately. In the highly diversified Coleoptera order, for instance, structural colouration has been intensively studied in certain families, such as in the Scarabaeidae, that display marked iridescent colours, or the Chrysomelidae. Vigneron et al. (18) investigated the vivid and iridescent blue colouration of *Hoplia coerulea* (Drury) (Coleoptera: Scarabaeidae) and attributed it to the layer-by-layer structure of the beetle cuticular squamule, that act like natural photonic devices. The same authors (19) studied the tortoise beetle *Charidotella egregia* (Boheman) (Coleoptera: Chrysomelidae), which is able to modify its structural colour under stress conditions thanks to humidity content variation in a chirped (*i.e.* showing a progressive decrease of the layers thickness) multilayer reflector. Another chirped multilayer reflector producing gold colouration was observed by Pasteels et al. (20) in another species of the same genus, *C. ambita* (Champion). Although these examples testify of structural based colouration research in Coleoptera, such studies have been neglected in some Coleopteran families, where pigments have received most attention. This is the case of the Coccinellidae family, commonly known as ladybirds or ladybugs, a highly diversified family with 6000 described species (21) which harbour a wide range of colours and patterns but whose colouration has been exclusively considered from the perspective of its pigmentary origin.

Melanins and carotenoids are pigments considered to underlie the iconic red with black spots colouration of many species of ladybirds (22). Melanins are phenolic biopolymers synthetized by animals, plants, fungus and even some bacteria (23). In insects two kinds of melanin are produced: the brown / black eumelanin and in some cases the reddish pheomelanin, that contribute in particular to cuticle sclerotization and colour (24). Ladybirds are supposed to only bear eumelanin (25). Carotenoids, on the other side, are terpenoids of yellow, orange and red colour and animals acquire them via their diet, with few exceptions (26–28). Trophic relationships in the Coccinellidae family are highly diverse, ranging from their well-known predatory habits to herbivory or mycophagy, and theoretically all these food sources can provide carotenoids to their consumers. However, studying the evolutionary ecology of such a highly colour polymorphic family exclusively based on the dichotomy carotenoids-melanins might be simplistic. Indeed, different physiological mechanisms and developmental or genetic pathways, including at nano-structural level, may be involved in the production of final colour phenotypes (16,29). The variation in ladybird colouration, ranging from beige to black and encompassing all shades of red-orange, may partly explain why pigments have received greater attention in the literature thus far. Furthermore, it is noteworthy that the most studied structural colours are green, blue or metallic (e.g. silver or gold), while red hues are more commonly attributed to pigments. However, Justyn *et al*. (9) have highlighted that, although non-iridescent red structural colour is rarely observed in nature (30,31), structural iridescent red has been documented (32). This type of colouration is often combined with pigments, which reduce iridescence by absorbing other visible wavelengths. Ladybirds’ elytra are hardened fore wings and have the typical laminar structure of insect cuticle, which is essentially composed of structural proteins and chitin but also contains water and in some cases pigments and metals (33–36). Concerning ladybirds’ elytra, their mechanical properties have been investigated by Zhou et al. for *Coccinella septempunctata L.* (37), but studies on ladybirds’ elytra are not extensive and, to our knowledge, there is no research work on the structural colouration of ladybirds.

Taking all the above into consideration, we defend that investigating the origin of colour and the evolution of colour-related interactions in ladybirds requires a broader perspective that includes not only pigments but structure as well. Therefore, we herein hypothesise that the colours of the Coccinellidae involve the contribution of both pigments and structures. To test this hypothesis, we will study the origin of colour in two species, *Adalia bipunctata* (L.) and *Calvia quatuordecimguttata* (L.), which are different in terms of elytra colour and patterns (Fig 1).

**Fig 1.**
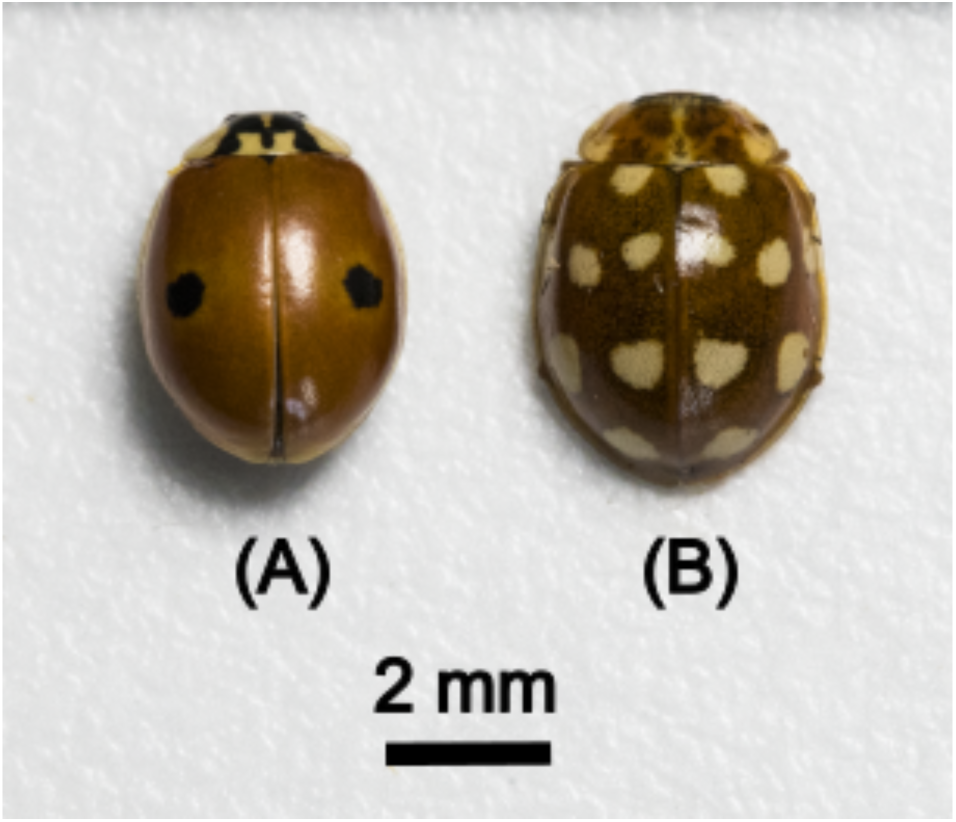
Photographs of an individual of both the studied species. Colouration and patterns of adults of (**A**) *Adalia bipunctata* (L), “typica” morph and (**B**) *Calvia quatuordecimguttata* (L).

*A. bipunctata* is polymorphic regarding to elytra pattern, and presents forms ranging from red to almost completely black elytra (25) (in the present study we analysed individuals from the “typica” morph; Fig 1(A). *C. quatordecimguttata,* on the other side, is monomorphic and the elytra appear brown with cream spots (Fig 1(B)), the colouration of which cannot be solely attributed to pigmentation as no known pigment is capable of producing the glossy white / creamy hue observed, thereby supporting the hypothesis that both structural and pigmentary factors contribute to the overall colour. Despite their phenotypic differences, the two species belong to the same tribe, the Coccinellini (21), are of Paleartic origin (38), are present in the same habitats, and commonly co-occur in communities of aphidophagous Coccinellidae (39). That is, they thrive in similar light and temperature contexts. Moreover, in the laboratory they can be reared exclusively on the same aphid species *Acyrthosiphon pisum* (Harris) (40,41) enabling control over their carotenoid’s food source.

Understanding how structure and pigments interact and contribute to the observed colouration in ladybirds, requires investigating both their physical and chemical properties. In the present work, several methods were used, ranging from optical and electron microscopies, light-based techniques, to LC-HRMS analyses.

## Material and methods

### Ladybird culture

Individuals of the two species were obtained from stock cultures maintained at the “Centre de Recherche sur la Biodiversité et l’Environnement” Laboratory (Toulouse, France). These consisted of adults reared at 21 ± 1°C, LD 16:8, in 5-litre plastic boxes with a piece of corrugated filter paper, on which the females laid eggs. Three times a week they were fed an excess of pea aphids, *<u>Acyrthosiphon</u> <u>pisum</u>* Harris. Two stems of broad bean, *Vicia faba* L., were added to each box to improve the survival of the aphids. Eggs were taken from the stock cultures every rearing day and incubated in 175 cm^3^ plastic boxes kept under the same conditions as described above. After hatching, larvae were fed excess pea aphids three times a week and reared to adult stage.

Adults studied in this work were between 15 and 20 days old and have developed their natural colouration. They were collected from the stock cultures, frozen and kept at - 21°C until analyses, in order to avoid the degradation at room temperature and air ambient. Just before the analyses, the elytra were removed with the help of a scalpel.

### Physical properties

#### Optical and electron microscopies

The elytra were analysed both at the surface and cross section levels. The elytra surface was first observed at the micron meter scale by a Nikon Eclipse LV100ND optical microscope in reflection mode and then analysed using Scanning Electron Microscopy (SEM), at nanometric scale, with a Helios NanoLab600i SEM. Other than allowing to image the surface at bigger magnification than optical microscopy, SEM detects morphology independently from light interaction with the samples.

In parallel, Transmission Electron Microscopy (TEM) was used in cross sections of elytra in order to investigate the elytra microstructure up to nanometric scale, and to identify the phenomena that could be responsible for their physical colour. Cross-sectional samples for TEM were prepared in CMEAB (Centre de Microscopie Electronique Appliquée à la Biologie, Toulouse, France). The sample preparation protocol differed between the two species. In the case of *C. quatuordecimguttata,* the elytra were fixed in 2% glutaraldehyde in 0.1 M Sorensen phosphate buffer (pH 7.4) for 4 hours at 4°C, followed by a 12-hour wash in 0.2 M phosphate buffer at 4°C. The samples were then post-fixed for 1 hour at room temperature in 1% osmium tetroxide with 250 mM sucrose and 0.05 M phosphate buffer. Dehydration was carried out through a graded ethanol series, followed by propylene oxide treatment, before embedding in Embed 812 resin (Electron Microscopy Sciences). In contrast, *A. bipunctata* elytra were directly embedded in the same resin, bypassing the preparatory steps required for *C. quatuordecimguttata* elytra, where direct resin embedding had proven ineffective. Polymerisation of the resin blocks was achieved over 48 hours at 60°C.

Finally, the elytra were sectioned into 70 nm-thick slices using an Ultracut Reichert Jung ultramicrotome fitted with a diamond knife and mounted on 100-mesh collodion-coated copper grids.

SEM and TEM images were analysed using the software ImageJ (54), version 2.14.0/1.54f.

#### Light based methods

Local measurements for reflectance spectra were performed in order to quantify the elytron colouration and investigate the light interaction with the elytra surface. Raman spectroscopy was used as a non-destructive method to detect the presence of pigments in the elytra. This is a physical method to detect chemical molecules vibration energies based on the inelastic scattering of light by molecules and has been already employed to characterize pigments (carotenoids, melanin…) present in different animal tissues (55–57). For these measurements, a X-Plora Horiba/Jobin Yvon device was used with a confocal microscope, a white light lamp, a 532 nm laser wavelength, and a 100X objective lens. Several gratings are available with 2400, 1800, 600 and 300 gr/mm (grooves per mm). Increasing the number of grooves/mm increases spectral resolution but reduces the observed spectral range and the signal intensity, so the right configuration has to be chosen on a case-by-case basis. In the present study, the 300 gr/mm grating was chosen for white light experiments (*i.e.* reflectance spectra recording), in order to obtain the larger spectral range and the 1800 gr/mm was used for Raman spectroscopy. To acquire reflectance spectra for the elytra, the reflected spectra registered in the direction of the microscope axis (normal reflection) was divided by a reference spectrum, corresponding to a perfect mirror in the visible range (100% reflector).

For all the analyses, 10 elytra from 10 different individuals of each species were studied. All the spectra were recorded in the same area of the elytra, close to the black or creamy spots depending on the species, and with the laser beam perpendicular to the elytron surface.

#### Numerical Simulations

Numerical simulations were conducted using the finite-difference time-domain (FDTD) method, implemented with the Meep software package (58). This electromagnetic simulation tool solves Maxwell’s equations iteratively over time within a finite computational region. The FDTD approach was chosen due to its accuracy in predicting the optical response in inhomogeneous multilayer structures (59). In particular, the thicknesses of each interlayer were adjusted to replicate the contrast alternation observed in the TEM cross-sectional images.

The computational grid resolution was set to 100 pixels per μm, with an additional subpixel smoothing routine. Absorbing boundary conditions were employed at the edges of the computational domain to minimize reflection artefacts. These conditions were achieved using a perfectly matched layer (PML) setup, which surrounds the computational cell with a non-reflective medium that absorbs outgoing electromagnetic waves.

To explore the optical properties of the system, we set the optical birefringence between layers to 0.15, while scanning the mean refractive index between 1.30 and 1.55, without any absorption (k=0). The incident light was generated from point sources randomly distributed in front of the structure, simulating a wide range of incidence angles. The broadband, incoherent electromagnetic wave source was designed to emulate natural daylight conditions. Reflectance spectra were obtained as the average of ten independent computational runs, each with slight random variations (with a standard deviation of 5 nm) in the geometry to account for inherent variability in the biological structure.

For chromaticity response analysis, each reflectance spectrum was multiplied by Planck’s spectral energy luminance function at 6000K in order to obtain the spectral radiance of the system under white light illumination. Then, we employed the CIE 1931 colour matching functions, which use a set of tristimulus values that correspond to the spectral sensitivities of the three types of cone cells in the human eye. This approach allowed for an accurate assessment of the perceived colour response of the simulated structures under daylight conditions (see supplementary material S1 for colour reproduction of a photographic standard colour chart).

### Chemical analyses

#### Sample preparation and extraction

Ladybug elytra samples (6 replicates, 2g each) were extracted using a solvent mixture of ethanol:hexane:MTBE (1:0.5:0.5). Samples were homogenized using a FastPrep-24 instrument (3 cycles of 20s at 4 ms−1, with 5-min cooling periods), then centrifuged (15 min, 3500 rpm, 4°C). The supernatants were collected and stored at 4°C. A quality control sample was created by pooling aliquots from all extracts for LC-HRMS analysis.

#### UHPLC–HRMS profiling, Peak Analysis, Features Identification, Statistical Analysis

See supplementary material S2

## Results

*A. bipunctata* elytra appear orange-reddish with black spots, while those of *C. quatuordecimguttata* appear brown with creamy spots. To human eye, iridescence is missing for both species. Fig 2 illustrates the reflectance spectra and the associated colours patches obtained for (A) the orange zone of *A. bipunctata* elytra and the creamy (B) and brown (C) zones of *C. quatuordecimguttata* elytra, respectively.

**Fig 2.**
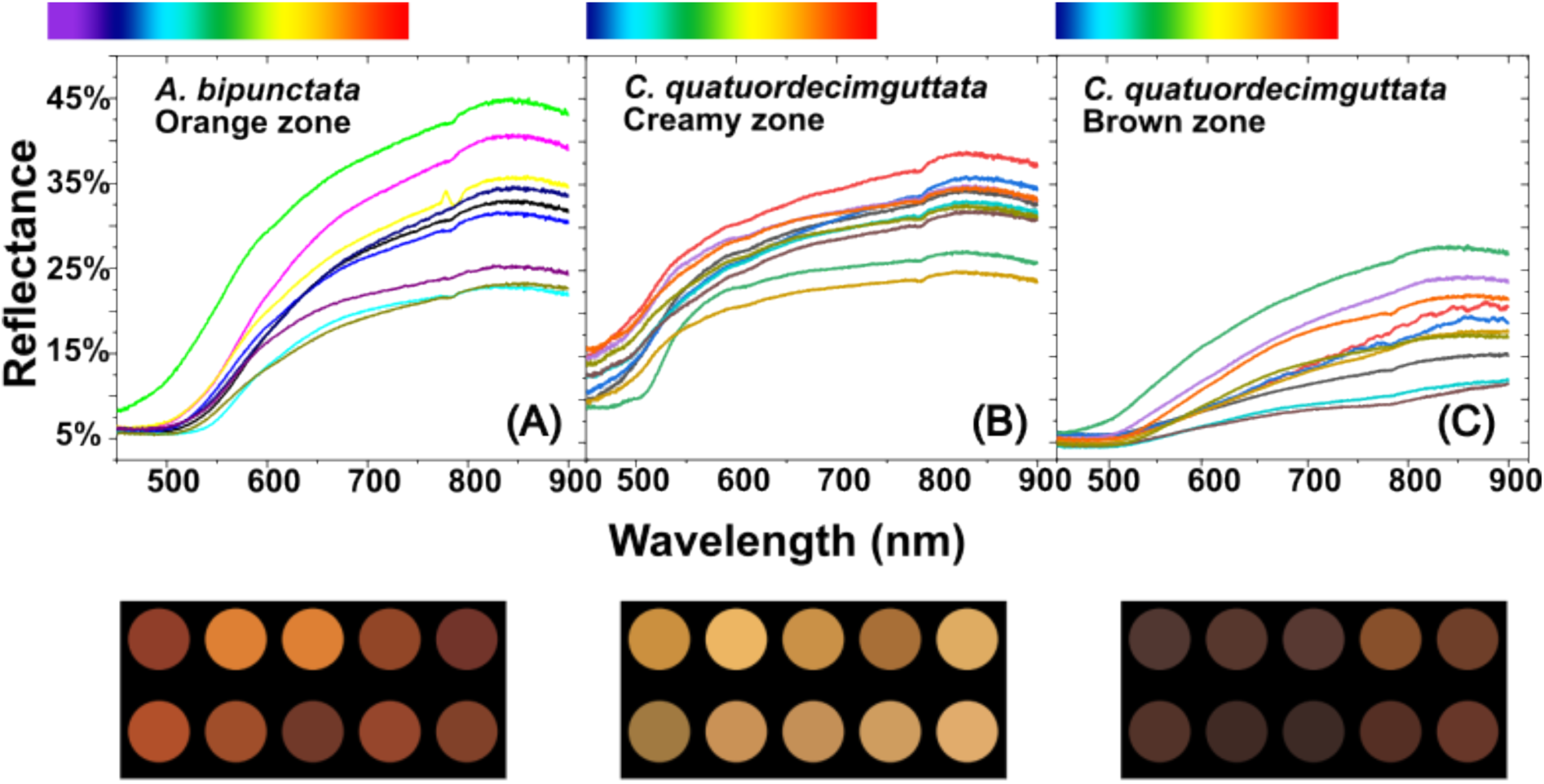
Reflectance spectra. The curves are the reflectance spectra of ten analysed individuals for each species and obtained for (**A**) the orange zone of *A. bipunctata* elytra and the creamy (**B**) and brown (**C**) zones of *C. quatuordecimguttata* elytra. A (human) colour scale corresponding to wavelengths axis is placed over each spectrum. The colour patches at the bottom of each figure correspond to the colours that are perceived for each of the reflectance spectra.

Reflectance spectra for the black spots of *A. bipunctata* are not shown due to the extremely low signal detected. For all the others observed elytra zones, light is predominantly reflected within the yellow-to-red range although maximum reflectance occurs in the infrared, which is invisible to human beings, insects, and birds (42) (a colour scale corresponding to wavelengths perceived by humans is added to each figure for clarity). Thus, the lowest reflectance values for the three observed zones are between 400 and 500 nm, which corresponds to the spectral zone where carotenoids typically absorb light (43). The perceived colour can be determined as described in the section “Numerical Simulations” by multiplying each reflectance spectrum by Planck’s spectral energy luminance function at 6000K and using the CIE 1931 colour matching functions. This results in the coloured patches at the bottom of the Figs 2A – 2C, which correspond to the ten reflectance spectra recorded from each elytra zone. Variations between individuals are observed in both chromaticity and luminance, with luminance referring to the light intensity, which is associated with the amount of reflected light. It is important to note that the measurements were taken at a highly localised scale (using a x100 lens), although the numerical aperture of the lens allows for light collection across a broad angle. Consequently, the coloured patches do not represent a uniform colouration across the entire elytron, and variations may occur even within a single elytron.

When the elytra of the two species are observed with the optical microscope in reflection mode, small-scale differences are detected in their morphology and colouration (Fig 3).

**Fig 3.**
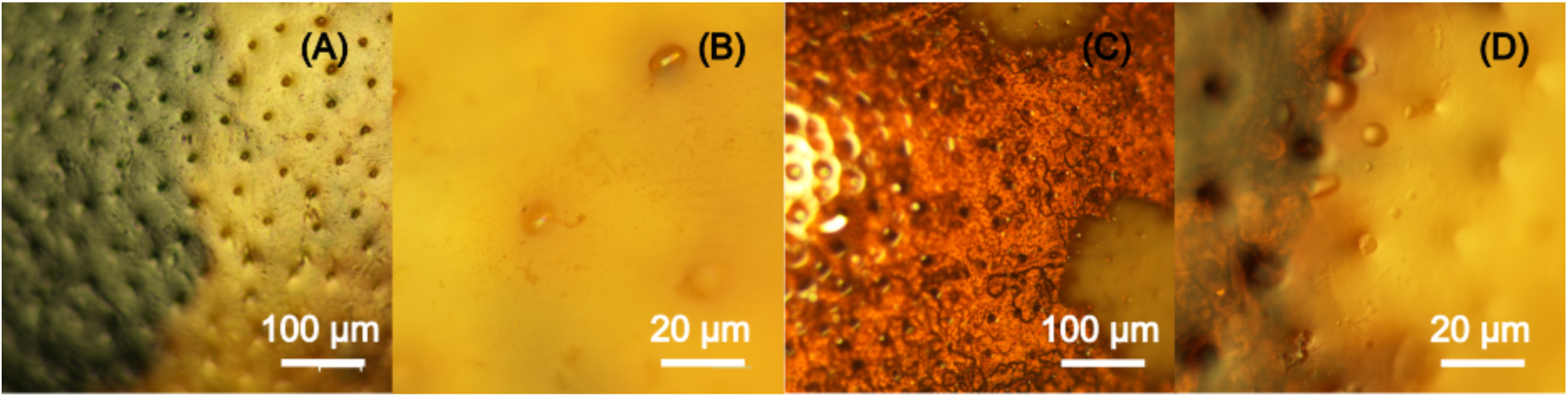
Optical microscopy. Optical microscope images of a same zone of the elytron surface of an *A. bipunctata* specimen (**A** and **B**) and of a *C. quatuordecimguttata* specimen (**C** and **D**).

*A. bipunctata* elytra present a rather regular surface colour, bearing concavities evenly distributed in the orange and black zones (Figs 3A and 3B). Contrarily, the colour of *C. quatuordecimguttata* elytra’s surface is not regular (Fig 3C). Whereas it is uniformly coloured in the bright/creamy spot zones, the brown zones are formed by red-brown and darker grains. As for *A. bipunctata*, concavities are also present and evenly distributed in the brown and creamy zones (Fig 3D). These cavities have a diameter of more than 5 17m and cannot be involved in colour production, as evidenced by the fact that, in both species, the zones between them are orange, black, brown, or creamy depending on the elytron zone.

The Scanning Electron Microscopy (SEM) images of the two species’ elytra reveal that concavities are in both cases seta-bearing (Fig 4A and 4C) and also show the existence of smaller concavities in between the larger ones (Figs 4B and 4D).

**Fig 4.**
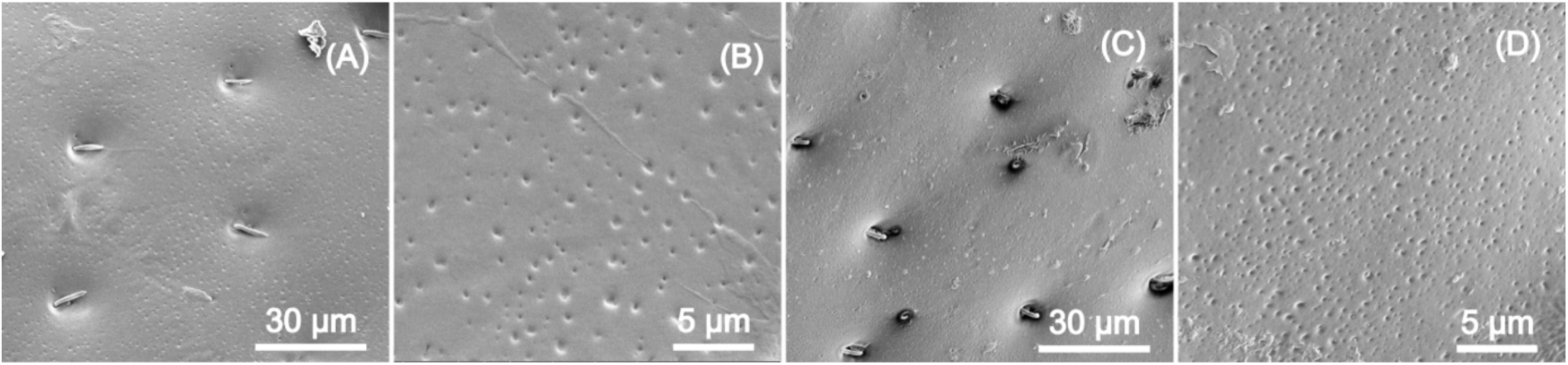
Scanning Electron Microscopy. SEM images of the surface of an *A. bipunctata* (**A** and **B**) and *of a C. quatuordecimguttata* elytra (**C** and **D**).

These smaller cavities have a diameter between 125 and 370 nm, and 75 and 400 nm respectively for *A. bipunctata* and *C. quatuordecimguttata.* Using the Euclidean Distance Measurements (EDM) approach in the Fiji (ImageJ) program, the mean distance between these smaller concavities is 814 nm ± 440 nm for the first species whereas it is 173 nm ± 100 nm for the second one.

Fig 5 depicts the Transmission electron microscopy (TEM) cross-section microstructure of the elytra of the two species.

**Fig 5.**
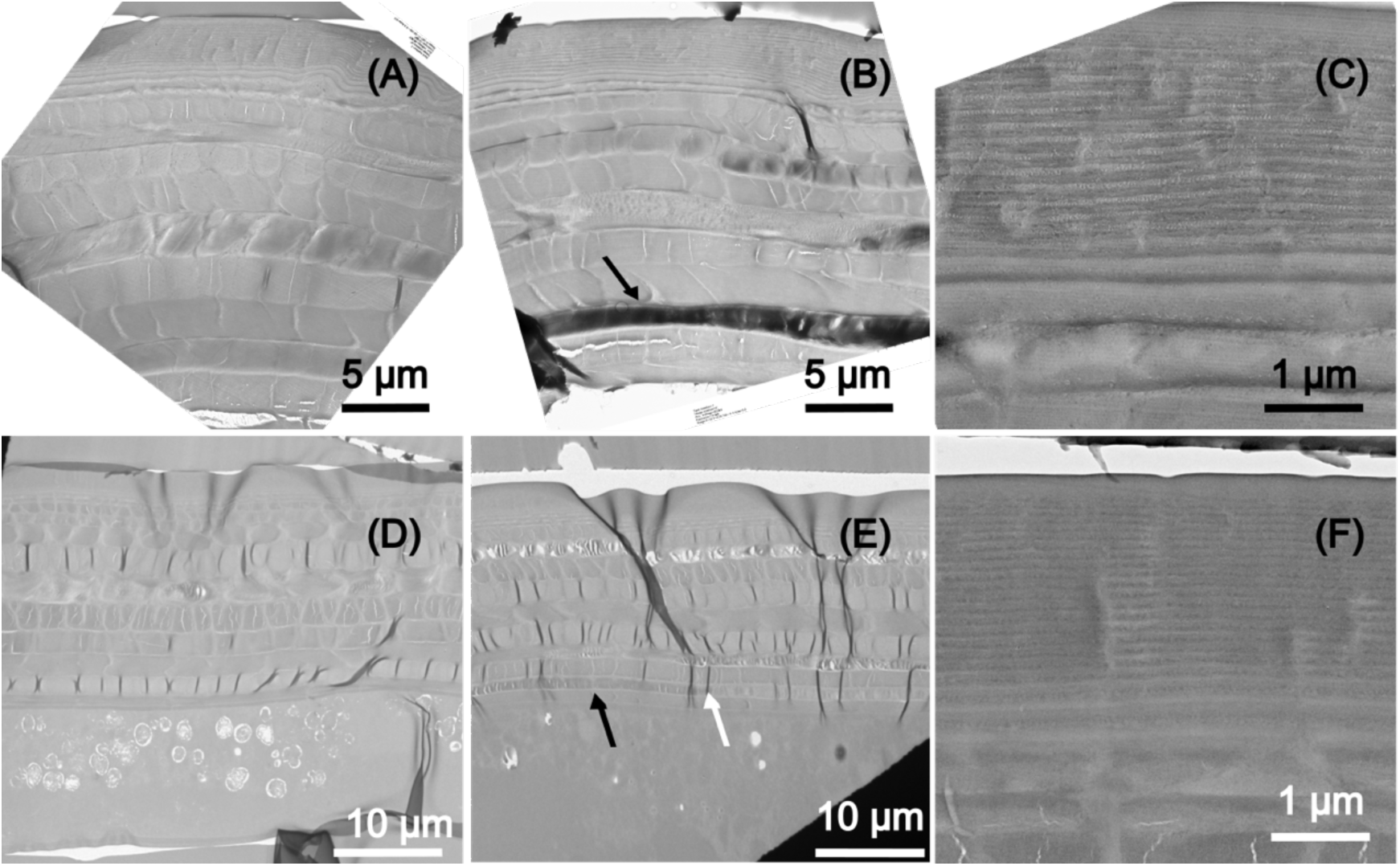
Transmission electron microscopy. TEM images of the cross-section microstructure of (top raw) an elytron of *A. bipunctata (***A**) section of an orange zone, (**B**) section of a black spot zone, (**C**) zoom on the exocuticle multilayer, and of (bottom raw) an elytron of *C. quatuordecimguttata (***D**) brown zone, (**E**) transition from brown and creamy spot zone, (**F**) exocuticle multilayer. Black arrows in panels B and E indicate the location of an electron high absorbing layer. The white arrow in panel E indicates the location of the frontier between the zones with and without the electron high absorbing molecule.

In both species the endocuticle presents a laminar structure made up of multilayers of darker and brighter contrast (Figs 5A and 5D). In the case of *A. bipunctata* these layers are 1 to 2 17m thick whereas in *C. quatuordecimguttata* they range from 1 to 2.2 17m thick. It should be noted that the cross-section microstructure of a black spot zone of the *A. bipunctata* elytra (Fig 5B) shows an electron-high absorbing layer (black arrow), indicating the presence of a molecule that is opaque to electrons, potentially eumelanin. A similar yet less contrasted layer is observed in the brown but not creamy zones of the *C. quatuordecimguttata* elytra (see black arrow in Fig 5E). Finally, the epidermis of *C. quatuordecimguttata* elytra is larger than that of the *A. bipunctata* elytra. The exocuticle of *A. bipunctata* examined at higher magnification corresponds to a band made of 30 thin layers, with a period spanning from 112 to 270 nanometres, and it has the same properties (total thickness and periods) on the orange and black zones. Furthermore, this exocuticle multilayer reveals the presence of several nanoparticles (30 to 40 nm diameter) within the thin layers (see Fig 5C). The exocuticle has been found to have the same structure for all the observed specimens. The exocuticle of *C. quatuordecimguttata* is also organised in a chirped multilayered structure, but it is made of 42 layers with periods varying from 103 nm to 270 nm. Nanoparticles are also visible in this layer, with 25-40 nm diameter (Fig 5F) and porosities of the same order of magnitude can be observed. In this species the exocuticle also has the same properties (total thickness and periods) on the brown and creamy zones, as shown in the Fig 5E, depicting the frontier (white arrow) between the brown zone of the elytron and a creamy spot.

The interaction of light with the chirped multilayer in the exocuticle of ladybirds has been modelled in accordance with the methodology outlined in the section “Numerical Simulations”. Fig 6 presents the simulated structural chromaticity, excluding the effects of light absorption from pigments and/or the matrix.

**Fig. 6:**
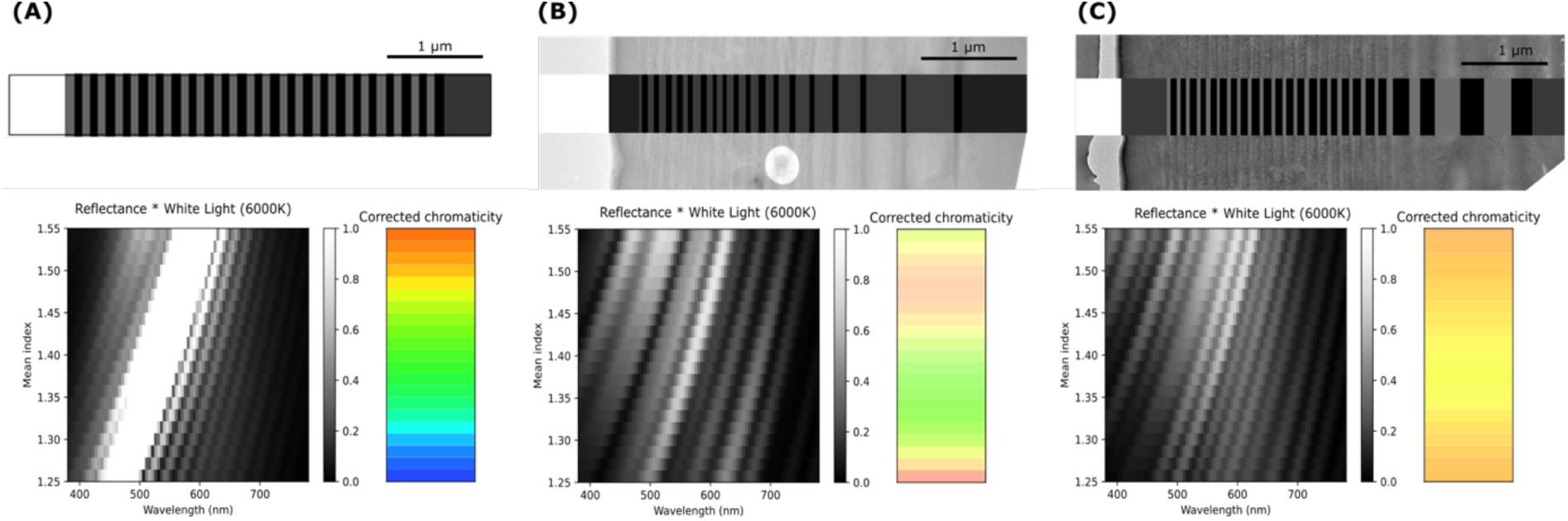
Simulated structural chromaticity. Top row: (**A**) Bragg grating with a fixed periodicity p=190 nmp=190nm; (**B**) Correspondence between the TEM cross-sectional image of the *A. bipunctata* elytra structure and its numerical replica; (**C**) Similar analysis for *C. quatuordecimguttata*. The grayscale represents variations in the refractive index. Bottom row: Dependence of reflection radiance on the mean refractive index and the corresponding CIE 1931 chromaticity coordinates.

Initially, a model Bragg grating with a fixed periodicity approximating that observed in the ladybird multilayers (p=190 nm, n=190 nm) was evaluated. This analysis demonstrated that such a structure is capable of selectively reflecting colour across the visible light spectrum, contingent on the mean refraction index. Subsequently, the numerical replica of the structures corresponding to the TEM cross-sectional images of the *A. bipunctata* (B) and *C. quatuordecimguttata* (C) elytra were analysed. The dependence of reflection radiance on the mean refractive index was calculated for each species, and the corresponding CIE 1931 chromaticity coordinates were determined. These simulations suggest that the structural characteristics of *A. bipunctata* would give rise to a colouration ranging from green to orange-pink, whereas those of *C. quatuordecimguttata* would produce a colouration that ranges from yellow to orange. The observed colouration is primarily determined by the average refractive index of the chitin matrix. However, the presence of pigments may lead to the absorption of certain reflected wavelengths, thereby influencing the final perceived colour. We therefore examined the pigments present in the elytra.

Fig 7 shows an analysis of elytra pigment content using Raman spectroscopy for the two species.

**Fig 7.**
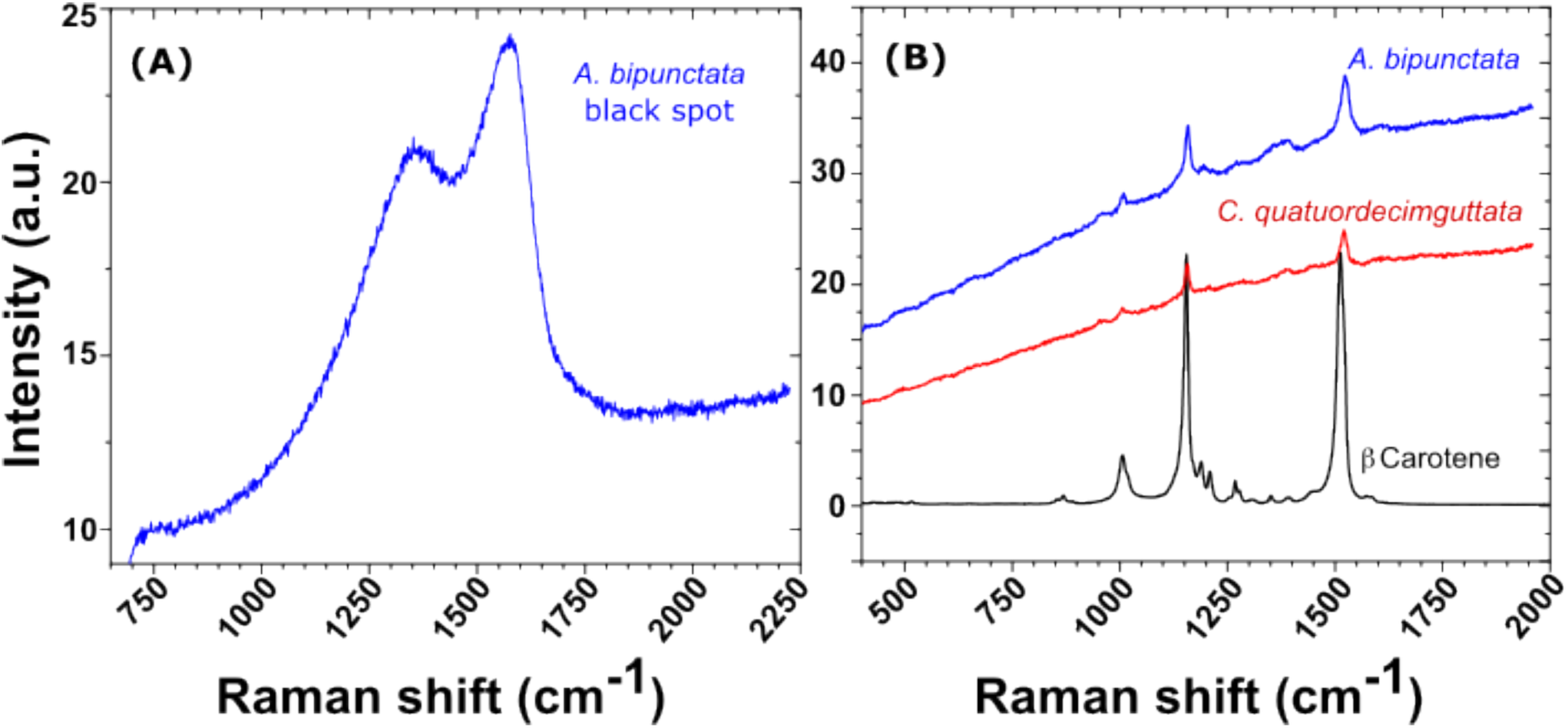
Raman spectra in different elytral zones. (A) Raman spectrum of the black dot of an *A. bipunctata* elytron. (B) superposition of beta carotene reference spectra with the Raman spectra obtained in the orange zone of an *A. bipunctata* and in the brown zone of a *C. quatuordecimguttata* elytra.

Fig 7A depicts a typical Raman spectrum obtained from the black dots on the elytra of *A. bipunctata* ladybirds. The spectrum structure and peak positions correlate to/match the reported Raman peaks for eumelanin, which are located at 500 cm^-1^, 1380 cm^-1^, and 1580 cm^-1^ (44). Because the Raman peaks’ positions are determined by the molecule’s surroundings, these values may vary slightly depending on the matrix. In our case the peaks are centred at 1380 cm^-1^ and 1567 cm^-1^. Raman peaks corresponding to eumelanin have not been identified in any additional regions of the elytra, including neither the orange area of *A. bipunctata* nor the brown areas of *C. quatuordecimguttata*. In Fig 7B, the Raman spectra obtained in the orange zone of an *A. bipunctata* elytron and in the brown zone of a *C. quatuordecimguttata* elytron are superposed with beta carotene reference spectra. The highest peaks in all of the spectra correspond to carotenoids. Indeed, the largest contributions of carotenoids to Raman spectra are in the ranges [1000; 1008] cm^-1^ (C-CH3 bonds), [1151; 1156] cm^-1^ (C-C bonds), and [1511; 1529] cm^-1^ (C = C bonds), suggesting the existence of these pigments in the elytra of both species. A study of Raman spectra from ten different individuals of each species confirms that carotenoids are present in both (Fig 8).

**Fig 8.**
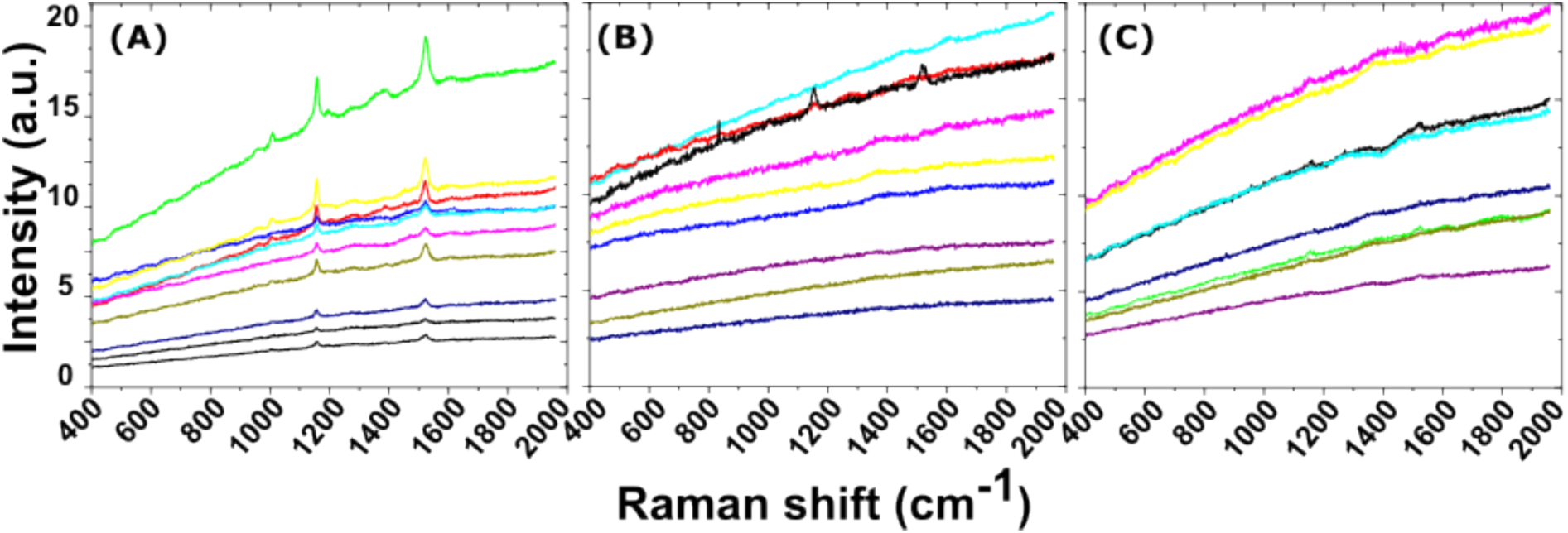
Variability in Raman Spectra. Raman spectra of the elytra of ten specimens of (**A**) *A. bipunctata* orange zone, and of ten specimens *C. quatuordecimguttata* (**B**) creamy and (**C**) brown zones

However, there is also intraspecific variation within each species. In the case of *A. bipunctata*, Raman spectra (Fig 8A) show very strong carotenoids’ signal, present in all individuals. In *C. quatuordecimguttata,* in contrast, carotenoids signal is very weak and detectable only in 3 out of the 10 analysed individuals, implying that carotenoid concentration can be very low or even absent in some individuals of this species. Interestingly, in the case of *C. quatuordecimguttata*, when carotenoids are present, the corresponding peaks appear in both the brown and creamy zones of the elytron.

The metabolomic analysis revealed two clearly distinct clusters corresponding to A. bipunctata and C. quatuordecimguttata samples (Fig 9), demonstrating a strong differentiation in their overall metabolic profiles.

**Fig 9.**
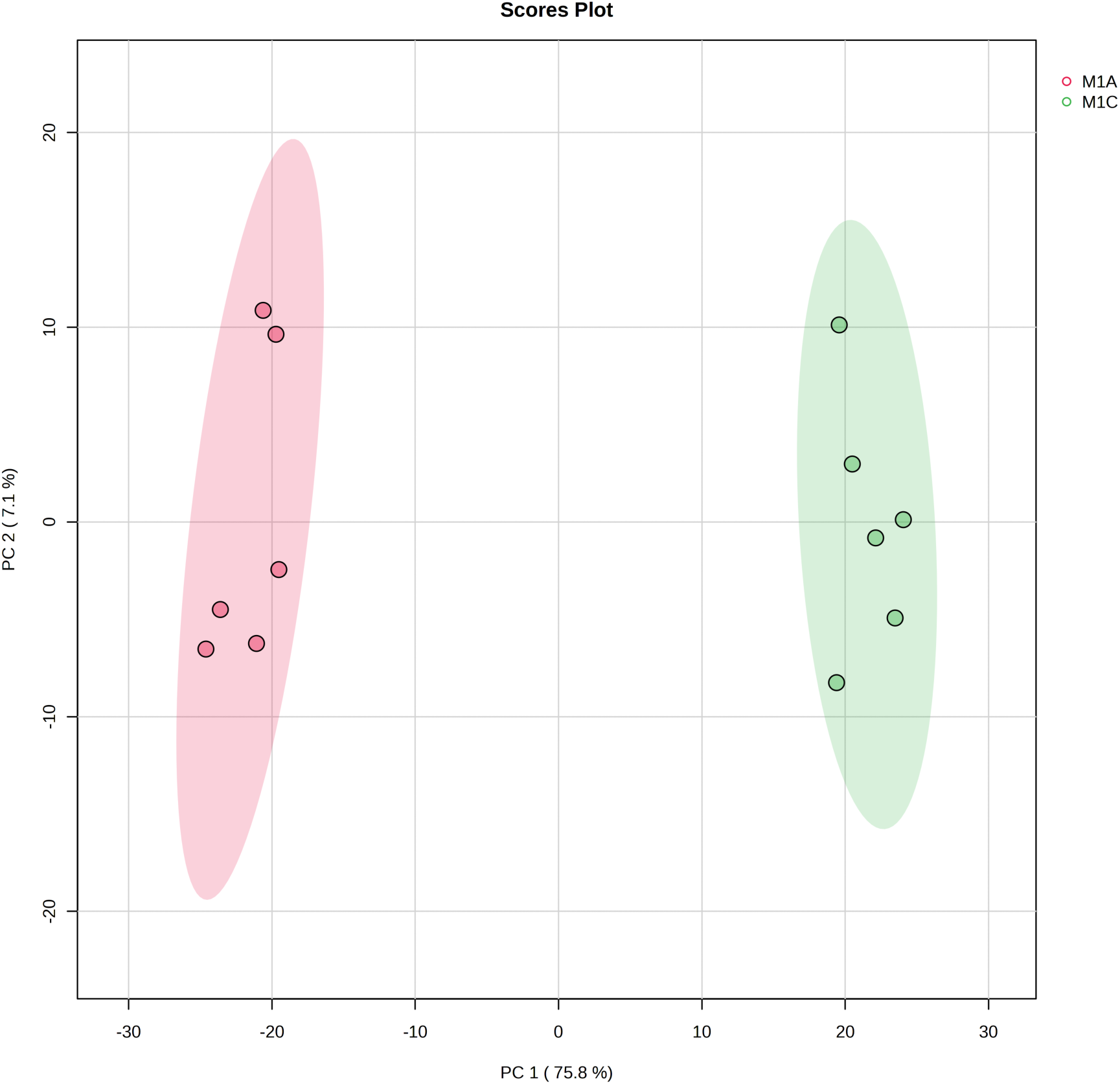
LC-MS Chromatograms. PCA corresponding to the extracts of *A. bipunctata and C. quatuordecimguttata* based on the variables (normalized peak area as a function of *m/z* at an Rt) detected on the LC-MS chromatograms.

When focusing specifically on pigment content, chemical analyses of the elytra revealed substantial differences in beta-carotene levels between the two species, confirming the Raman results. A prominent peak corresponding to beta-carotene (m/z 536.43750/Rt 11.413 minutes) was detected in *A. bipunctata*, whereas *C. quatuordecimguttata* samples showed minimal beta-carotene presence (data not shown).

## Discussion

The traditional approach to the study of animal colouration typically separates pigmentary and structural origins. However, we argue that this separation is potentially misleading. Pigments do not exist in isolation; they are invariably embedded within a matrix - the primary constituent of the tegument – which possesses intrinsic optical properties that both modulate the absorption characteristics of the pigments and are influenced by them in return. Structural colouration arises from the design of the tegument (such as its periodicity, size, and organisation) but it also depends on the material’s intrinsic properties, particularly its ability to refract light. This last one is quantified by the refractive index *n* which is a characteristic of the material and varies with wavelength. Importantly, the refractive index of a material also reflects the influence of pigments: when a range of wavelengths is absorbed, the refractive index becomes a complex number N = *n* + i*k*, where the imaginary component *k* accounts for light attenuation due to absorption. Thus, “physical” and “chemical” colouration are interrelated through the refractive index, with both pigments and the matrix material contributing to the overall optical properties of the tegument. Consequently, teguments should be considered as integrated optical media. The main question, therefore, is not whether the colouration is pigmentary or whether pigments are present, but how to appropriately assess their impact on the final colour, *i.e.* understanding how pigments affect the final colouration by attenuating specific wavelengths of light.

Given these considerations, we herein revisit the commonly held belief that the colouration of ladybirds’ elytra is due to pigments, particularly carotenoids and melanin. We focused on two species of ladybirds: *A. bipunctata* and *C. quatuordecimguttata*.

Our empirical results demonstrate that pigments, including carotenoids and/or melanin, are present in both species. These pigments may be embedded within the chitin of the entire elytron or reside in specific layers within the chitin matrix, as observed for the black spots of *A. bipunctata* and the brown zones of *C. quatuordecimguttata*, respectively. Indeed, the TEM analysis revealed the presence of layers with dark contrast in the endocuticle of these regions, which are absent in the red or creamy zones of the elytra. This darker contrast likely results from the presence of an electron-absorbing molecule, presumably a pigment, suggesting that differences in colours within an elytron are determined by pigments distribution. This supports our hypothesis that pigments modulate the structural colouration of the elytron. Notably, except for the presence of the dark contrast layer, the microstructure is consistent across all zones of the elytron (spots and ground colour) in each species, indicating that a single physical mechanism governs wavelength selection in each species.

The black spots of *A. bipunctata* are associated with the presence of eumelanin, as identified by Raman spectroscopy and producing the typical dark contrast in TEM cross sectional images. Eumelanin absorbs all wavelengths across the visible spectrum, irrespective of the structural selection of wavelength, making it the primary contributor to the black pigmentation. In contrast, we were unable to identify, by chemical analyses and Raman spectroscopy, the pigment producing the dark contrast in TEM images of *C. quatuordecimguttata* brown zones remains unidentified through both chemical analyses and Raman spectroscopy. A potential candidate is eumelanin, which may be present at a lower density compared to the black spots of *A. bipunctata*, thereby accounting for the reduced contrast. Alternatively, pheomelanin, a form of melanin that produces reddish-brown to yellow pigmentation, could be involved, as it is the case for certain insect species (45). However, its characteristic Raman peaks centred at 500, 1490, and 2000 cm⁻¹ (44) are conspicuously absent from the spectra obtained from all regions of the elytra of both species. Another possible candidate is a pigment from the pteridine family, responsible for yellow, orange or red colouration, which has recently been shown to be synthesised by insects (46). However, we didn’t observe any fluorescence under UV light, which is characteristic of pteridines.

With regard to carotenoids, TEM did not reveal any specific layers with contrast variations indicative of carotenoid presence. High-magnification analysis of the exocuticle multilayer in both species revealed nanoparticles with diameters between 25 nm and 40 nm, showing a darker contrast than the surrounding matrix, suggesting they may represent pigment aggregates. These aggregates highlight the complex interplay between structural elements and pigments in colour production. Raman spectroscopy revealed variations in carotene peak intensities between the two species, with a stronger signal detected in *A. bipunctata* compared to *C. quatuordecimguttata*, suggesting a higher concentration of carotenoids in the former species, corroborating chemical analyses. Additionally, Raman peaks from both the brown and creamy regions of *C. quatuordecimguttata* elytra exhibited similar intensities, implying that carotenoids do not account for the colour differences between the spots and the remaining elytron. Notably, no known pigment is capable of producing the shiny white/cream colour of the spots in *C. quatuordecimguttata*, suggesting that this colouration is not pigment-based but rather results from the structural design of the elytron and the optical properties of its components.

We cannot exclude that other pigments are present in the elytra of both species. Indeed, the effectiveness of Raman detection may be curtailed, particularly when pigments are situated in deeper layers. Furthermore, Raman spectroscopy is particularly sensitive to molecules with a high number of non-polar bonds, such as carotenoids, while pigments with fewer non-polar bonds are less efficiently detected. As a result, certain pigments may evade detection if their concentrations fall below the Raman detection threshold or are masked by carotenoids in presence of multiple pigments.

However, if pigments are responsible for modulating colour across different zones of the elytra in the two studied species, the fundamental question remains: what are the physical mechanisms governing the interaction between light and the elytra structure, in conjunction with pigments, that produce the final perceived colour? It is important to note that light interaction with the elytra structure may involve different physical mechanisms, which must be considered prior to assessing the origin of colouration in ladybirds.

The two primary physical mechanisms responsible for colouration in biological tissues are incoherent scattering (or diffusion) and coherent scattering (*i.e.* interference or diffraction). The operation of these mechanisms within the elytral structure and the specific structural elements involved are described below.

Structural non-iridescent colouration generally arises from light diffusion by particles, a phenomenon whose theoretical treatment is formalised by Mie theory (47). This accounts for the weak diffusion of red wavelengths, explaining why red colouration is difficult to achieve through light diffusion alone: when scattering conditions favour red wavelengths, the diffusion of other visible wavelengths results in an overall white/creamy appearance. The scattering efficiency of different visible wavelengths is discussed in greater detail in (31). The smaller concavities observed on the surface of the elytra of the species studied here may incoherently scatter light, producing a white or cream colour as seen in the scales of *Pieris rapae* (L.) (Lepidoptera), where a creamy hue results from scattering by densely packed pterin granules (pterinosomes) (48). In these butterflies, the size of the isolated particles is insufficient for optimal scattering, but their close packing leads to dependent scattering producing light diffusion across the visible spectrum and resulting in a white/creamy hue. In the elytra of both ladybird species, the concavities range from 100 nm to 400 nm, similar to the size of *P. rapae* granules, and could contribute to colour production by scattering light across all wavelengths. However, the density of concavities is higher in *C. quatuordecimguttata* than in *A. bipunctata*, making dependent scattering more likely in the former species. As a result, *C. quatuordecimguttata* is expected to exhibit more intense scattering effects compared to *A. bipunctata*, where this effect can be considered negligible.

In contrast with the previous diffusion mechanism, structural colours produced by coherent scattering depend on the optical path length travelled by light and vary with the observer’s angle, typically resulting in iridescence. Iridescence is not observed in the ladybird elytra examined here, but the presence of the observed pigments within the cuticle matrix may suppress iridescence by absorbing certain reflected wavelengths. As a consequence, the absence of iridescence does not necessarily rule out the generation of interferential colours. Cross sectional analysis of the elytra shows, for both species, a laminar structure characteristic of beetle elytra, with alternating dark and bright regions visible in TEM images. The endocuticle sublayers in both species are at least one micrometre thick, which is too thick to generate interferential colours within the visible range. However, the exocuticle in both species contains a chirped (i.e., with variable thickness) multilayer that we have shown to function as a reflector in the visible range, as it has already been observed in other beetles, notably in *Charidotella egregia* (Boheman) (19). Indeed, our numerical simulations indicate for both the investigated ladybird’s species that the light interaction with these structures produces wavelength selection. The chirped multilayers in the exocuticle are interferential layers that are able to generate a structural colouration whose chromaticity depends on the average refractive index of the chitin matrix.

The elytral structure of ladybirds is complex and the light interactions that induce colour production can include surface scattering, interference from multilayers and absorption from embedded pigments. As a result, significant challenges exist in predicting perceived colour. One of such challenges is the need to estimate not only the real part of the refractive index but also the imaginary part, which is related to the level of pigment absorption. The primary challenge when predicting the colouration produced by an interferential multilayer, such as those in the ladybird exocuticle, lies in determining the average refractive index and the refractive indices of each sub-layer. While spectroscopic ellipsometry has been successfully employed to obtain complex refractive indices in other beetle species (49), conducting similar measurements on ladybird elytra is exceptionally difficult due to their morphology, as the curved surface distorts the elliptical shape of the beam, complicating the measurements. Previous studies on the colouration of beetle bodies or elytra have used average refractive indices for chitin to estimate the wavelength of maximum reflection. Indeed, although approximations are necessary, when the period is relatively constant, it is possible to roughly estimate the colour produced by multilayers, as suggested by Vigneron *et al.* (19). For their calculations, these authors used the refractive index most commonly cited for chitin in recent literature, which is 1.56. However, this value may decrease depending on the water and air content within the chitin matrix. Furthermore, in the present study, which deals with chirped layers of increasing thickness, there is no simple method to determine the reflected colour, and numerical simulations were employed here to predict the full spectrum of reflected light.

The results of our simulations are presented as a function of the refractive index to address the problem of approximating its value. Moreover, by accounting for variations in the refractive index, we can simultaneously consider colour variations induced by density, air, and water in the matrix.

In this study, only the real part of the refractive index was considered, with the imaginary part assumed to be zero. To evaluate the imaginary component of the refractive index (i.e. the absorption contribution), future work will need to provide a more comprehensive understanding of the pigments present in the elytra and their concentrations. Our preliminary chemical investigations provide complementary insights into the pigmentary component of this complex system. The significant difference in beta-carotene content between *A. bipunctata* and *C. quatuordecimguttata*, alongside the clear metabolomic discrimination between these species, opens new perspectives for investigating the chemical basis of both matrix and pigments in Coccinellidae. This approach could reveal how pigment composition interacts with structural features to produce the final colour phenotype, potentially identifying new compounds involved in this interplay. Future studies combining metabolomics with structural analysis could provide a more comprehensive understanding of colour production in these insects.

However, our results already demonstrate that not only the interferential layers produce structural colouration, but that the chromatic response differs between species. It is evident that the chromaticity produced by the interferential layer alone does not correspond to the perceived colour of the elytra. However, if we consider the presence of pigments that lead to the absorption of a range of reflected wavelengths and/or additional phenomena such as scattering, the origin of the colouration in each of the species studied can be elucidated. Based on our results, we propose the following scenario to explain the origin of colouration in ladybirds:

In the elytra of A. bipunctata, coherent scattering from the chirped multilayer structure appears to be the main mechanism for physical colour production. The resulting chromaticity changes from green to orange in accordance with the refractive index value. Specifically, orange hues are obtained for refractive index values between 1.45 and 1.50. When the refractive index is 1.55, which is very close to the value commonly used for chitin, the colour is green. The final perceived colouration is influenced by pigments: carotenoids in the ground colour of the elytron and melanin pigments in the black spots. For the ground colour, when the refractive index induces a green structural colour, which is within the carotenoid maximum absorption range, an orange hue is produced. If the refractive index already favours an orange colour, the presence of carotenoids enhances this hue, potentially leading to a more intense colouration. This finding is particularly interesting for ladybird ecology, as it suggests that, for a given carotenoid content in the elytra, the perceived brightness or intensity of the colour is influenced by the refractive index of the structure, which in turn is determined by its density and composition, particularly its protein and water content. Regarding the black spots, as previously mentioned, eumelanin absorbs all wavelengths across the visible spectrum, irrespective of the structural wavelength selection, and is responsible for their black appearance.

For *C. quatuordecimguttata*, the hue produced by the chirped multilayer in the endocuticle remains within the light yellow to orange range. The chromatic response obtained when n=1.3 appears to be the closest to the creamy aspect of the spots. However, as discussed earlier, scattering resulting from the high density of surface cavities is also a possible explanation for this hue, and both mechanisms may coexist. Even though carotenoids have been detected in very low concentrations in both the creamy and brown regions of the elytra, they are not present in all the analysed individuals, suggesting that they do not contribute to the perceived colour. Regardless of the specific scattering mechanism, the brown ground colouration in this species may be associated with the presence of an as-yet unidentified pigment that absorbs light within the visible spectrum.

To conclude, we show here that the structure and not only the pigments contribute to the cuticle’s hue. Therefore, the study of ladybird’s colouration must consider the interplay between both elements. For instance, untapping the genetic basis of ladybird’s colouration will also require the study of the genes and molecular basis influencing the structural arrangement of the cuticle additionally to those associated with pigments (50). More importantly, understanding the evolution of colouration could benefit from considering structure. For example, the intensity of a colour patch or its area is expected to be a function of carotenoid deposition and thus of the entailed costs, which supposedly ensures the honesty of colouration as a signal of an individual’s quality (2,3). However, as structure influences colouration and, consequently, the signal sent to receivers (*e.g.*, potential predators), it may finally also affect signal honesty. In fact this may explain why carotenoids and melanin fall short of explaining that the intra and interspecific relative toxicity of ladybirds with contrasting colours do not consistently align with their degree of warning colouration (51–53). Considering that the structure of the cuticle contributes to colouration may open avenues to better understand how toxicity and colourations are linked. Our results, which demonstrate the interplay between chemical and structural colouration of the cuticle pave the way to a more comprehensive study of colour signalling in this model system.

## Supporting information

S1-colour-chart-reproduction

S2-Experimental Section_Chemistry

## Acknowledgments

The author would like to thank Dominique Goudounèche and Isabelle Fourquaux of CMEAB (Centre de Microscopie Electronique Appliquée à la Biologie) for electron microscopy sample preparation and assistance in SEM and TEM observations.

## Notes

### Competing Interest Statement

The authors have declared no competing interest.

