## Supplementary material for "Decoding ladybird’s colours: structural mechanisms of colour production and pigment modulation": S1-colour-chart-reproduction

### S1-Colour reproduction of a photographic standard colour chart

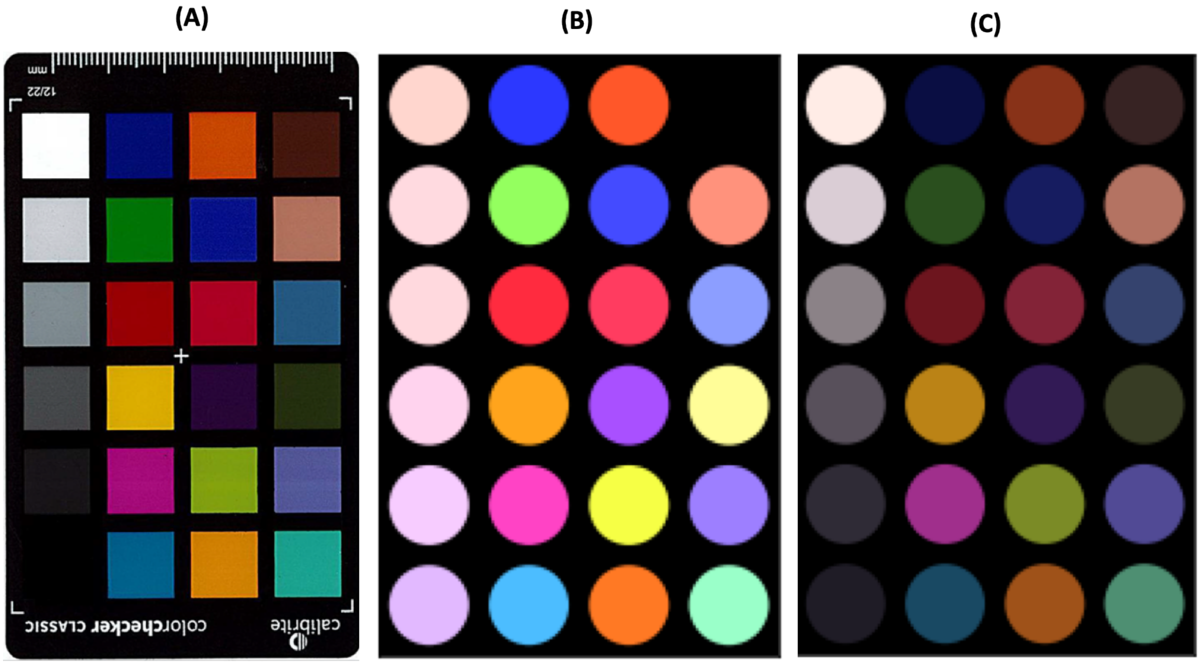


The reflectance spectrum of each colour in the photographic colour chart (Calibrite Color Checker) (A) was recorded under the same experimental conditions as those used for the ladybird elytra. (B) was obtained by directly applying the CIE 1931 colour matching functions to the reflectance spectra. (C) The reflectance spectra were multiplied by Planck's spectral energy luminance function at 6000K to obtain the spectral radiance of the system under white light illumination. The CIE 1931 colour matching functions were then applied.
