## Supplementary material for "Decoding ladybird’s colours: structural mechanisms of colour production and pigment modulation": S2-Experimental Section_Chemistry

***UHPLC–HRMS profiling***

All extracts were profiled using a UHPLC-DAD-LTQ Orbitrap XL instrument (Ultimate 3000, Thermo Fisher Scientific, Hemel Hempstead, UK). Two µl of the sample (final concentration of 2 mg/ml) were loaded onto a C18 Acquity column (100 × 2.1 mm i.d., 1.7 µm, Waters, MA, USA) equipped with a guard column. The analysis was performed using Waters system at a flow rate of 0.3 mL/ min. The following gradient was employed using water/0.1% formic acid (solvent A) and acetonitrile/0.1% formic acid (solvent B): time 0 min, 95 % A; 12 min, 95 % B. The instrument settings were as follows: Mass detection was performed using an electro- spray source in positive ionization (PI) and negative ionization (NI) modes at 15,000 resolving power (full width at half maximum (FWHM) at 400 *m/z*). The mass scanning range was *m/z* 100 – 1500 Da. The capillary temperature was 300°C and IS pray voltage was fixed at 4.2 kV (positive mode) and 3.0 kV (negative mode). The mass measurement was externally calibrated before starting the experiment. Each full MS scan was followed by data dependent MS/MS on the three most intense peaks using stepped collision-induced dissociation (CID) (35% normalized collision energy, isolation width 2 Da, activation Q 0.250).

***Peak Analysis***

The UHPLC/HR-MS raw data were processed with MS- DIAL version 2.70. Automatic feature detection was performed between 0.3 and 13 min for mass signal extraction between 100 and 1500 Da in positive and negative mode. MS1 and MS2 tolerance were set to 0.01 and 0.4 Da, respectively, in centroid mode. The resulting peak list was then exported to Microsoft Excel. Peaks were retained in the sample if appeared in at least 4 out of 6 replicates. The matrix was exported to comma-separated value (CSV) format prior analysis using MetaboAnalyst 3.0 with normalization using Pareto scaling for Two-tailed Student's *t*-tests to test the significance of differences between two species.

***Significant Features Identification***

The molecular formula and structure of significant features were calculated with MS-FINDER-RIKEN PRiMe version 2.28. A score is attributed to each putative compound considering bond dissociation energies, mass accuracies, fragment linkages and nine hydrogen rearrangement rules. Various parameters were used to reduce the number of potential candidates, such as the element selection exclusively including C, H, O; mass tolerance fixed to 10 ppm and the isotopic ratio tolerance set to 20%. All components were searched against the home database using SMILES input from the Dictionary of Natural product (CRC Press v26 : 2) specific for the fungal database, SciFinder, and also a built-in database in MS-FINDER system: Natural Products Database (UNPD), KNApSAcK, CheBI (Chemical Entities of Biological Interest), STOFF, T3DB (the toxin and toxin target database), NANPDB (Northern African Natural Products Database), DrugBank, FooDB, and PlantCyc. The compound molecular formula was retained for identified features with a score above 7 and only structures with a score above 5 were retained for thorough analysis.

***Statistical Analysis***

Statistical analysis was done by uploading the.*csv* file to the Metaboanalyst platform version 5.0 (https:// www.metaboanalyst.ca/). The data were pre-processed, first by normalization (by median) of the samples, then the variables were weighted by auto scale (mean-centered and divided by the square root of the standard deviation of each variable). A Principal Component Analysis (PCA) then a Partial Least Squares Discriminant Analysis (PLS-DA) were carried out. Finally, the *m/z* and the retention time of the significant features associated with their normalized peak area were plotted on a heat map (hierarchical clustering) using ANOVA. To test the significance of differences between two-group data (*Adalia vs Calvia*) t-tests ($p$-value < 0.05) were performed.
